# Cell-sheet shape transformation by internally-driven, oriented forces

**DOI:** 10.1101/2024.11.28.625908

**Authors:** Junrou Huang, Juan Chen, Yimin Luo

## Abstract

During morphogenesis, cells collectively execute directional forces that drive programmed folding and growth of the layers, forming tissues and organs. The ability to recapitulate aspects of these processes *in vitro* would constitute a significant leap forward in the field of tissue engineering. Free-standing, self-organizing, cell-laden matrices are fabricated using a sequential deposition approach that uses liquid crystal-templated hydrogel fibers to direct cell arrangements. The orientation of hydrogel fibers is controlled using flow or boundary cues, while their microstructures are controlled by depletion interaction and probed by scattering and microscopy. These fibers effectively direct cells embedded in a collagen matrix, creating multilayer structures through contact guidance and by leveraging steric interactions amongst the cells. In uniformly aligned cell matrices, oriented cells exert traction forces that can induce preferential contraction of the matrix. Simultaneously, the matrix densifies and develops anisotropy through cell remodeling. Such an approach can be extended to create cell arrangements with arbitrary in-plane patterns, allowing for coordinated cell forces and pre-programmed, macroscopic shape changes. This work reveals a fundamentally new path for controlled force generation, emphasizing the role of a carefully designed initial orientational field for manipulating shape transformations of reconstituted matrices.

## 1 Introduction

Tissues undergo complex morphogenesis during their development, and their capacity to execute these processes is encoded, in part, in their anisotropic cellular assemblies. The relationship between the extracellular matrix (ECM) and cellular activities is largely reciprocal. Anisotropy in the ECM provides a cue to directional cell proliferation and apoptosis, which are implicated in the elongation of drosophila embryos [1] and zebrafish gastrulation [2]. Anisotropic matrix mechanical properties induce gradients in cell behaviors. At the cellular level, cells undergo individual and collective durotaxis [3, 4, 5]. At the organism level, developing embryos exhibit a decreased stiffness and viscosity along the elongating posterior axis [6]. Conversely, cells such as fibroblasts are well-known to secret fibronectin and modify their environment [7, 8, 9]. As a result, anisotropy in ECM can be amplified through anisotropic division and gradual matrix remodeling. Increased anisotropy promotes further cell elongation and more efficient packing, prompting more division, thereby establishing a feedback loop [10]. This sequence of events eventually culminate in macroscopic shape changes. The ability to recapitulate and manipulate key aspects of these processes *in vitro* would constitute a significant leap forward in the field of tissue engineering.

Despite an abundance of strategies developed for creating hydrogels that can respond to stimuli such as light [11], heat [12], or pH [13], harnessing internally generated forces could greatly simplify experimental setup, while also better mirror the actuation strategies observed in nature. Recent years have witnessed a burgeoning interest in integrating living and non-living components to develop materials with spontaneous force generation and emergent properties. Internally driven systems have been realized by fueling an actomyosin network contractility with ATP in solution [14], by seeding fibroblasts on shape-changing polymer ribbon [15], by patterning mesenchymal condensates to program strain [16], and by harnessing yeast proliferation in hydrogels, in response to biochemical or physical stimuli [17]. Forces generated by cell traction can align matrix fibers [18], which help orientate neighboring cells [19]. The transmission of the force is coordinated across multiple cells, through parallel stress fibers organized at the site of cell-matrix adhesion for fibroblasts [20]. Below a threshold of cell-cell distances, cells permanently com-pact collagen matrices [7]. However, typically remodeling occurs isotropically; orchestrating collective cell forces would also require precise control over the directionality of the individual cells.

To resolve this issue, cell-laden collagen matrices can be aligned using mechanical stretching [21], but this approach lacks the necessary flexibility in the type of patterns that can be achieved. Aligned collagen matrices have also been generated by microfluidic devices [22], though the resulting geometrical patterns remain limited. One notable development in recent years has been the incorporation of fragmented electrospun fibers into 3D printing. These fibers align under flow conditions, thereby inducing the alignment of cells extruded alongside them [23, 24]. Nonetheless, 3D printing is constrained in the precision of the size of the nozzle, often with a lower limit of 100s of *µ*m. Hence, it will be difficult to accurately replicate regions experiencing large changes of orientation, for instance, near topological defects [25, 26]. These topological defects are hypothesized to function as organizational centers during tissue morphogenesis [27, 28], making them valuable structures to replicate *in vitro*. Furthermore, implementing these strategies requires specialized instrumentations typically not available in standard biological laboratories.

A growing body of literature draws parallels between cell ordering and liquid crystals. Steric interaction between elongated cells prompts them to align with their neighbors. In experiments, elongated cells follow boundaries similar to classic LC anchoring [29, 30, 31], their spatial arrangements engender topological defects [32, 33, 26], and they also spontaneously form patterns and undergo phase transitions [30, 34, 35]. Indeed, anisotropic cues in the form of nematic order is a straightforward way of directing cell assembly and regulating cell density [26]. Prepatterned liquid crystal elastomer (LCE) sheets are a versatile and powerful platform for driving programmable shape morphogenesis [36, 37]. This naturally raises the question of whether a similar strategy can be applied to a prepatterned 2D cell layer, by subsequently inducing transformation through intercellular forces. However, most cells are adherent, hence the internally generated stresses in the monolayer are tightly coupled to the substrate, making shape changes difficult to achieve. There have been attempts to remove the template after cells form the desired arrangement [38, 39], still, our ability to recapitulate alignment and morphogenesis remains limited.

In this paper, we demonstrate the fabrication of initially flat, thin, free-standing, cell-laden matrix sheets with controllable shape-morphing capabilities. First, we fabricate hydrogels with fibrin-like morphologies templated by a lyotropic chromonic liquid crystal, ordered by flow or boundary cues. Next, we characterize the anisotropic hydrogel microstructures by scattering and microscopy. Thereafter, we embed human dermal fibroblasts in collagen and grow them on pre-ordered hydrogels. Cells align following the anisotropic cues and remodel the collagen. These cell-laden collagen sheets spontaneously contract through the traction force of the encapsulated cells. By programming the orientation of the cells, we also control the macroscopic shape transformation of the matrix. We demonstrate that different cell orientations lead to distinct final shapes. These oriented cell sheets are modular, requiring minimal instrumentation and no special release mechanism, aiding their wide adaptation. This work establishes the groundwork for generating matrices that can undergo orientationally directed shape transformations, paving the way for the ultimate realization of *in vitro* morphogenesis.

## 2 Results

### 2.1 Controlling the microstructures of PEG hydrogels

Here, we fabricate polyethylene glycol (PEG) diacrylate (DA)-based hydrogels templated by a lyotropic chromonic liquid crystal (LC) disodium cromoglycate (DSCG), which form stack-like aggregates. The mixture of these two components forms a new LC phase. Previous research [40, 41] highlights the importance of chain length, concentration, and temperature conditions in stabilizing the nematic phase of the PEG-DSCG mixture, which is the origin of the fiber-like structures. Short-chain PEGDA minimally disrupts the LC phase and forms large structures with long-range order through its association with DSCG, despite lacking an LC phase on its own. In this work, we fix the composition (5 wt% PEGDA250 and 15 wt% DSCG) and focus on investigating different strategies to sculpt the architecture of the fibers. Importantly, the orientation of PEG fibers encodes the signal for directing cell alignment.

Each approach offers distinct advantages (Fig. 1a): (i) the rubbed substrate is the simplest to implement, whereas (ii) flow alignment in glass capillary tubes is preferred when a greater thickness is desired, and (iii) photopatterning is ideal for achieving spatially defined orientations. In all cases, similar fiber-like microstructures are obtained. Hydrogels, prepared in contact with surfaces that are either untreated or mechanically rubbed, were visualized using scanning electron microscopy (Fig. 1b-c(i)). Microstructures of PEG hydrogels prepared by flow alignment and photopatterning are shown in Fig. S1.

**Figure 1:**
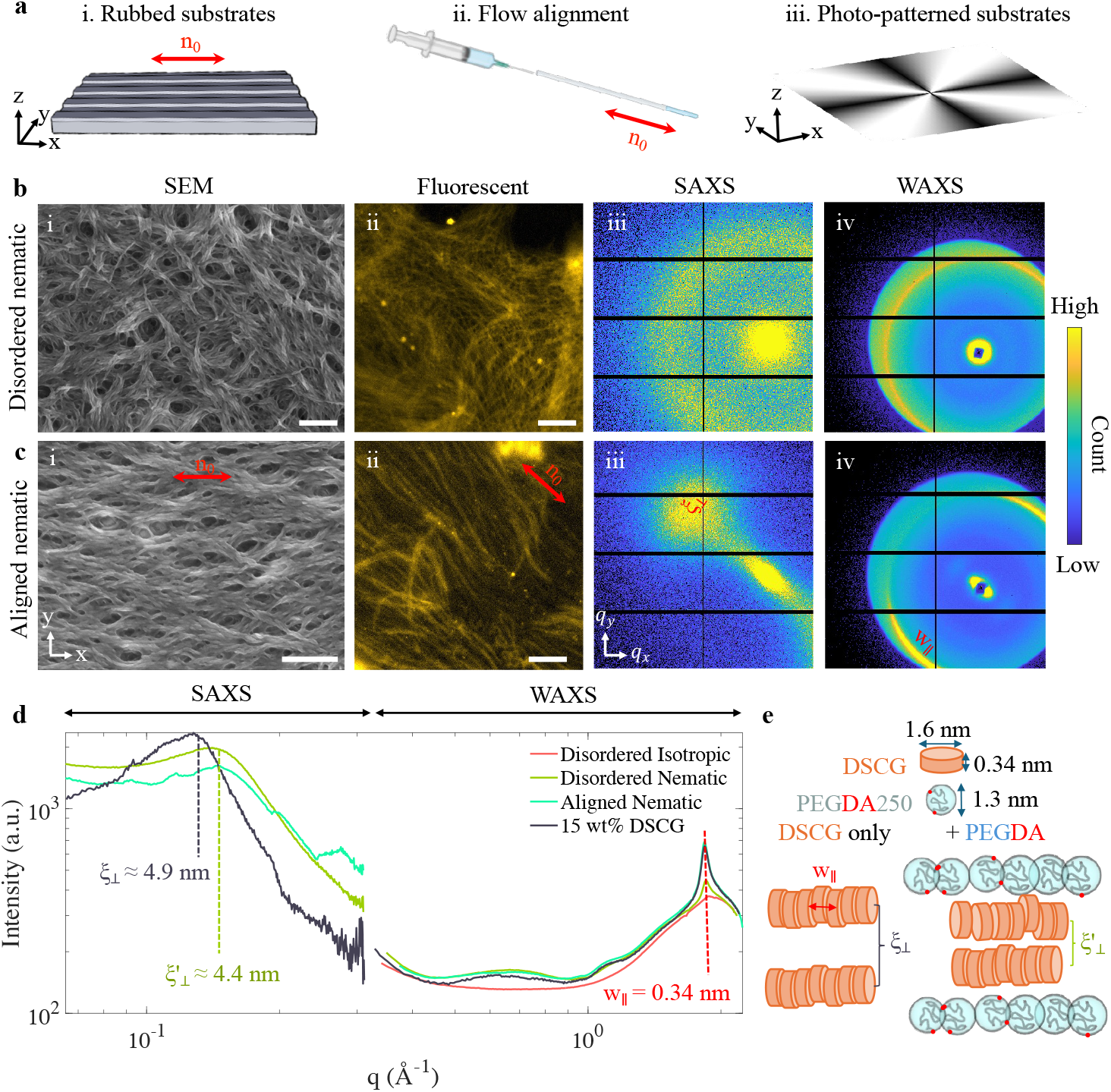
(a) Graphic representation of the three methods used in this study to generate hydrogel alignment. Double-sided arrows denote the alignment direction. (b-c) Microstructures of fibrous PEG hydrogels for (b) disordered nematic and (c) aligned nematic. (i) Air-dried structures, fabricated on substrate without or with rubbing, captured by scanning electron microscopy (SEM). The scale bars are 1 *µ*m. (ii) Aligned and disordered hydrogels tagged by rhodamine B, captured *in situ* by fluorescent microscopy. The scale bars are 50 *µ*m. (iii) Small-angle X-ray scattering (SAXS) and (iv) wide-angle X-ray scattering (WAXS). (d) Combined SAXS and WAXS scattering intensities for disordered and ordered PEG fibers, and references. (e) Schematics depicting the dimension of the PEGDA monomer and DSCG, and the postulated microscopic picture based on the scattering data.

To understand how the incorporation of PEGDA monomers modifies the DSCG microstructure *in situ*, we fluorescently label the hydrogels (Fig. 1b-c(ii)). Fluorescent microscopy indicates that individual PEG fibers are separated from their neighbors by vast space, as we use a wt% of PEGDA far below the overlap concentration of 34 wt%. Hence, PEGDA is depleted from the bulk and recruited to the DSCG surface, forming a “sheath”. This flexibility potentially enhances the interpenetration between PEG hydrogels and the collagen, facilitating cell contact guidance.

We further characterize the hydrogel microstructures by X-ray scattering (Fig. 1b-c(iii-iv)). To compensate for the low scattering contrast of the PEGDA in water, we prepare scattering samples in a 1.5 mm quartz tube by flow alignment to create a large scattering cross-section. Disordered sample, isotropic sample (15 wt% DSCG, 5 wt% PEGDA, crosslinked at T = 60 °C), and uncrosslinked pure 15 wt% DSCG are also prepared and scattered for reference. Crosslinking is performed at T = 5 °C for all the nematic samples, solidifying the molecular structure. Small- and wide-angle X-ray scattering (SAXS and WAXS) are shown for disordered samples in Fig. 1b(iii-iv), which are distinct from the anisotropic scattering patterns in Fig. 1c(iii-iv). The anisotropy is further highlighted in Fig. S2, which shows the intensity versus angle plot at wavevector q = 0.145 Å^−1^, with distinct peaks for the anisotropic sample. The merged SAXS and WAXS intensity plot is shown in Fig. 1d. We use subscripts “‖” and “⊥” to denote parallel and perpendicular to the alignment direction. In the WAXS regime, all samples, including the isotropic ones, exhibit a peak at q = 1.85 Å^−1^, which corresponds to the stacking distance of DSCG at 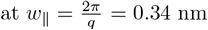, which is the thickness of a single DSCG molecule (Fig. 1e), consistent with earlier studies [42, 43].

The most notable differences appear in the WAXS regime, where a peak shift is noted from q = 0.13 Å^−1^ to 0.14 Å^−1^ for pure 15 wt% DSCG to all 5 wt% PEGDA, 15 wt% DSCG samples. DSCG naturally forms stack-like aggregates (Fig. 1e). Given that this peak is perpendicular to *w*_∥_ in the SAXS plot of the aligned nematic sample (Fig. 1c(iv)), we deduce that it must correspond to the distance between neighboring stacks of DSCG, *ξ*_⊥_ (Fig. 1c(iii)). The phase shift indicates that the inter-aggregate distance of DSCG decreases from *ξ*_⊥_ = 4.9 nm to 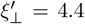 nm upon adding PEGDA. This is consistent with a microscopic picture where PEGDAs exert osmotic stress at low concentrations, which reduces the distance between aggregates [44]. This likely results from microphase separation and leads to the sheath-like structure. Note that with increased PEGDA chain length or concentration, the osmotic stress could eventually lead to a complete collapse of the inter-aggregate distance, resulting in macroscopic phase separation or phase change, consistent with prior observations [45]. In the main text, we present scattering results of hydrogels after crosslinking and washing off DSCG, consistent with the preparation used in the rest of the study. In Supporting Information (SI), we also present scattering results for samples immediately after crosslinking without washing off the DSCG and observe identical scattering patterns (Fig. S3). This has led us to infer that for crosslinked samples, the intensity represents the empty space left behind by DSCG upon washing, which has similar signals as DSCG which scatters much more strongly than the surrounding PEG. This indicates that PEGDA replicates the structure of the DSCG microstructure. Taken together, these results indicate that the liquid crystallinity of the PEGDA-DSCG mixture ensures its adherence to boundaries and flow sensitivity and establishes the hydrogels’ overall architecture, whereas the interaction between PEGDA and DSCG governs the microstructures.

### 2.2 Anisotropic contraction of the freestanding, cell-laden collagen sheet

These all-PEG hydrogel fibers resemble the fibrillar proteins that comprise the ECM. Inspired by how the fibrous proteins guide cell migration, we demonstrate the use of aligned PEG hydrogels as physical cues to direct cell alignment in a 3D environment. While 2D cell alignment on these hydrogels has been demonstrated [40], here, the alignment guidance at the boundary propagates through cell-cell contact.

This is first illustrated with uniformly aligned PEG hydrogels. To enable cell spreading into an extended morphology in 3D, the surrounding matrix must be degradable. Therefore, we select collagen, a natural polymer, to illustrate the alignment guidance of PEG hydrogel which can extend into a 3D volume. We deposit a suspension of neonatal human dermal fibroblasts (HDFs) suspended in neutralized Type I collagen onto aligned PEG fibers. Next, we place a glass coverslip on top of the collagen, separated by polydimethylsiloxane (PDMS) spacers (h ≈ 100 *µ*m), and then incubate the collagen-cells mixture at 37 °C to facilitate collagen crosslinking. After 1 hour, we release the coverslip. The cells are then cultured for 5 to 6 days. Afterward, the collagen layer readily separates from the hydrogel fibers underneath, and we obtain a free-standing cell-laden collagen sheet (Fig. 2a). The cell layer is several cells thick, as shown by confocal microscopy (Fig. 2b). Cells align along the tracks established by the PEG hydrogels, which act as “guide rails” through contact guidance between cells and the undulated topography. HDFs align along the x-direction, consistent with the hydrogel fiber orientation. Cell alignment is attributed to directional proliferation, collagen remodeling by the cells, and steric interactions amongst the cells. Furthermore, we have ruled out the possibility that this alignment is merely a result of topographical or mechanical effects by testing hydrophobic LCE fibers, which have a modulus of approximately 100 MPa. In this case, cells exhibit a much lower degree of alignment (S = 0.14, Fig. S4). While both LCE and PEG fibers can effectively guide cell alignment in 2D [46, 40], we find that this effect extends to 3D alignment only with the PEG hydrogel. Due to the unique softness and hydrophilic nature of the PEG hydrogel fibers, cells follow the directional guidance as they migrate through the collagen.

**Figure 2:**
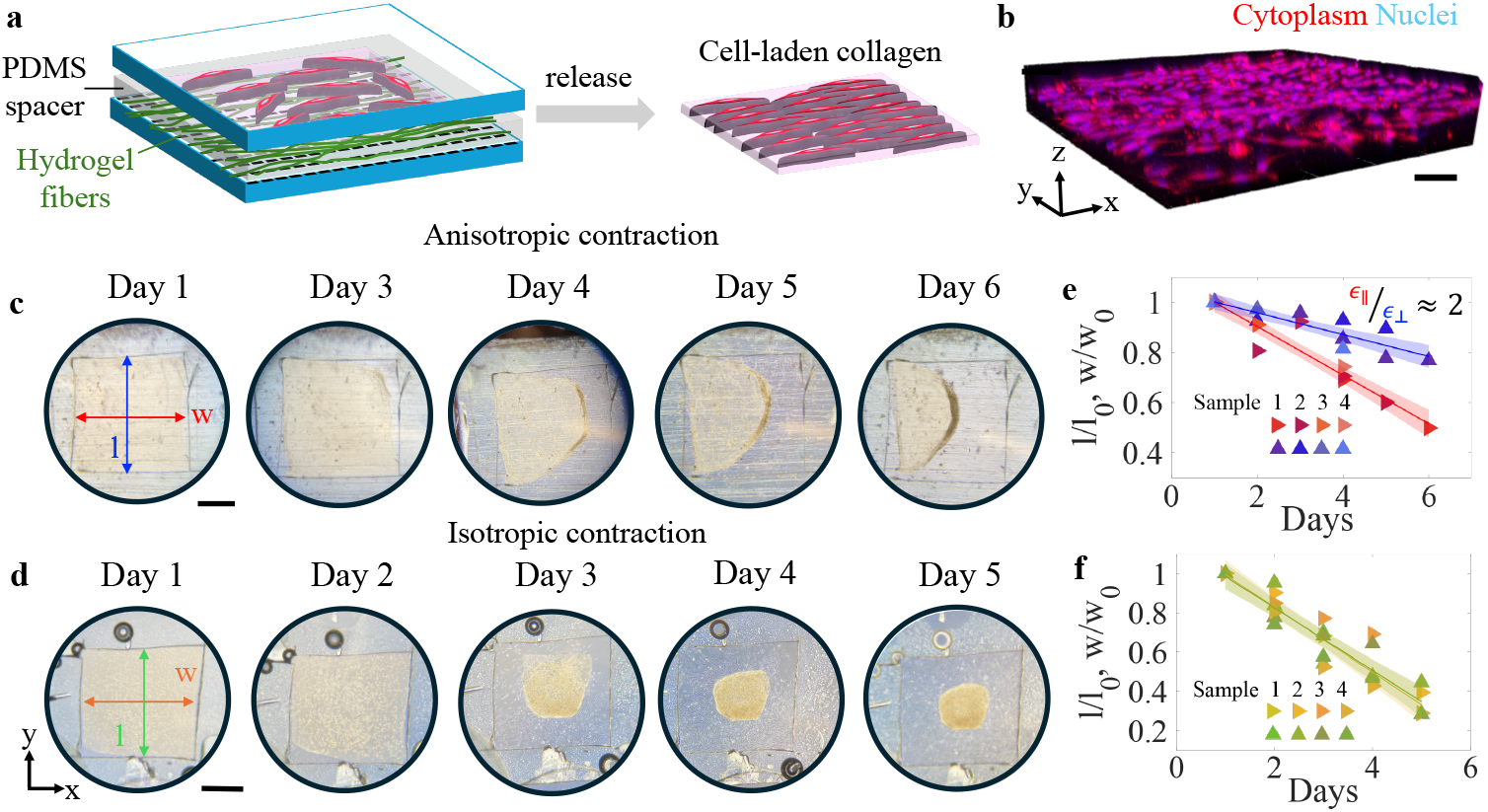
Anisotropic contraction of cell-laden collagen sheet. (a) The schematic depicts the fabrication of aligned cell assemblies. After removing the hydrogel template, cells remain aligned. (b) Confocal microscopy showing cell alignment. The scale bar is 100 *µ*m. (c) Macroscopic contraction of an initial 3 mm ×3 mm square of collagen encapsulates cells with alignment. The collagen contrasts anisotropically, more rapidly along the alignment direction, due to cell traction force. (d) In contrast, collagen with unaligned cells contracts isotropically. The scale bars are 1 mm. The region within the collagen gels is highlighted for clarity. (e-f) Width *w* and length *l* over time, normalized with respect to initial dimensions. Lines represent linear regression. Shaded regions indicate confidence interval from the fitting.

We postulate that the force exerted by thousands of cells, arranged in an anisotropic manner, collectively over time, can induce enough deformation which manifests in macroscopic anisotropic shape changes. This is demonstrated in Fig. 2c, using a free-standing collagen sheet with uniformly aligned cells similar to what is shown in Fig. 2b. The collagen sheet contracts significantly more along the alignment direction. In comparison, HDFs grown in hydrogel without alignment contract isotropically (Fig. 2d). HDFs are seeded at density *ρ* ≈ 2.5 × 10^5^ mL^−1^ in collagen and grown for several days. The cell growth rate is comparable within both anisotropic and disordered collagen matrices during this time - the cell number increased from 25 to 800 per mm^2^ in both isotropic and the aligned case, and the number density is uniform across both samples, which was determined by counting the number of cells in different regions of the substrate from the projection of the cells at a mid-height. These results indicate that the difference in contraction rate between the two samples arises from the orientation of the cells, rather than a difference in their distribution or growth rate.

To gain more quantitative insights into the relative rate of contraction, we document the contraction daily and use a custom MATLAB script to extract the width *w*, and length *l* of the collagen sheet (n = 4) by inputting 5-7 points along each edge. The procedure is illustrated in Fig. S5. For anisotropic samples, we perform a linear regression using all data and obtained two slopes 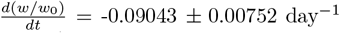, whereas 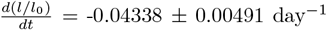, or a contraction ratio 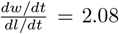 (Fig. 2e), which means that a typical sheet with aligned cells contracts approximately twice as quickly along the alignment direction. For isotropic samples, we only observe one slope from linear regression, thus we conclude that the contraction rate are the same in perpendicular and horizontal direction (Fig. 2f). Across all samples tested (n = 4), HDFs embedded in collagen in contact with the aligned hydrogels show a slower collagen contraction, which we attribute to the time required for them to remodel the network — an effect we will illustrate in the next section. Previous studies have noted the ability of collective cell behavior to compact freely suspended collagen gels [47]. Here, we show that we can harness the coordinated action of thousands of cells to induce millimeter-scale deformation.

### 2.3 Modification of the collagen microstructure

The collagen layer functions not merely as a passive scaffold but also hosts essential cell activities. The relationship is reciprocal: HDFs develop order due to their contact with the hydrogel fibers. On the other hand, fibroblasts can permanently compact collagen matrices [7]. As a result, culturing the cells inside the hydrogel also modifies its structure (Fig. 3a,b), thus cell-ECM interactions also beget anisotropic mechanical properties. The progression of cell alignment is tracked by fluorescent microscopy images taken on Day 0 and Day 5 (Fig. 3c,d). We extract the cell orientation by fitting an ellipse to the nucleus of each cell, where *θ* denotes the orientation of the long axis with respect to the alignment direction. The polar histograms of *θ* are shown in Fig. 3e,f. Cell alignment is quantified by 2D order parameter *S* = ⟨2 cos^2^ *θ* _−_ 1⟩, where ⟨·⟩ ensemble average for n ≈ 2200 cells. Fig. 3e corresponds to *S* = 0.053, with almost no order, whereas Fig. 3f corresponds to *S* = 0.48, exhibiting a high degree of alignment.

**Figure 3:**
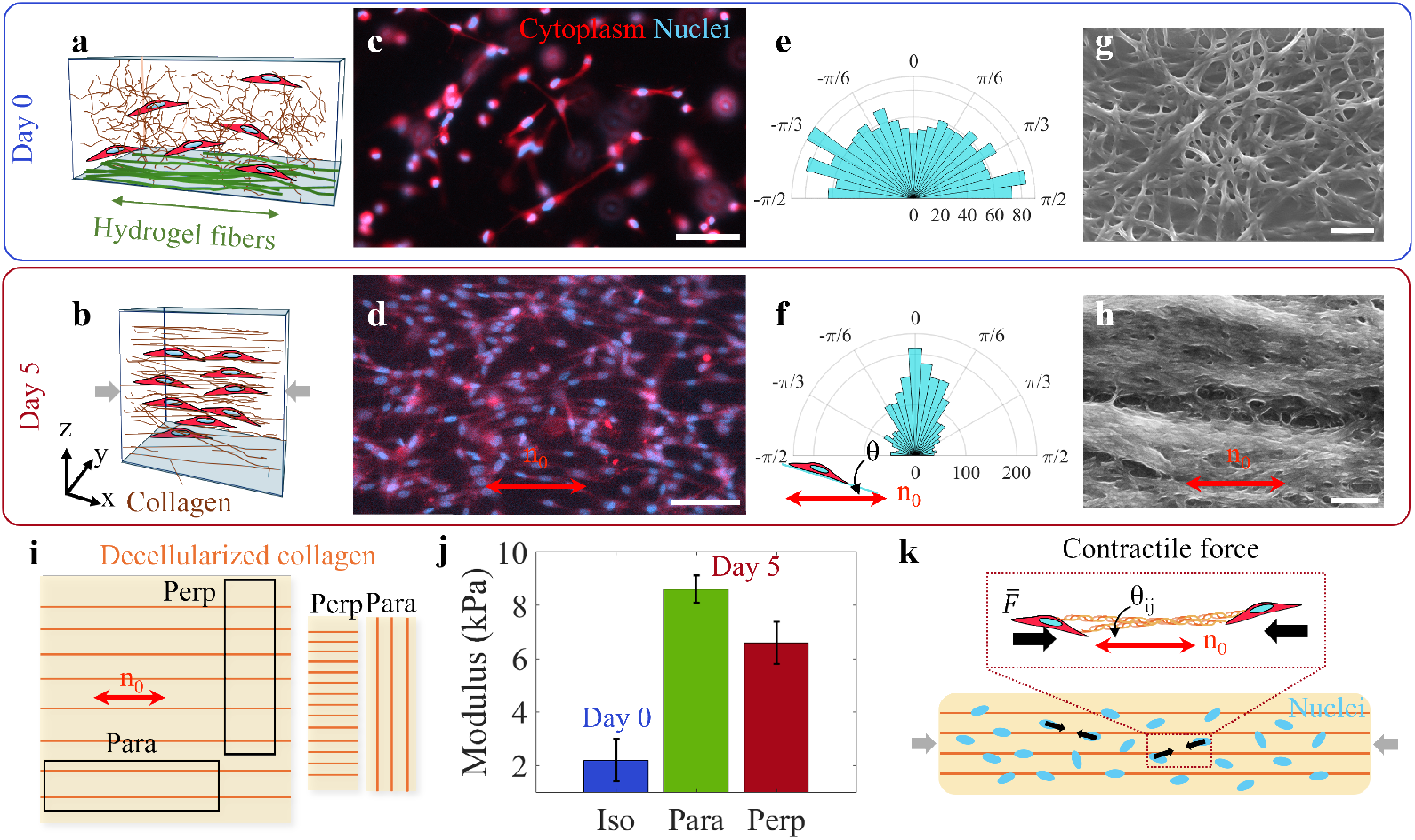
Cell alignment and modification of the collagen matrix at two time points (Day 0 and Day 5) during the contraction. (a-b) show schematics depicting the interaction between cells and the matrix. (c-d) Fluorescent micrographs of cell morphologies. The scale bars are 100 *µ*m. (e-f) The orientations of the cells are plotted in the polar histograms. (g-h) Scanning electron micrographs of the collagen. The scale bars are 1 *µ*m. (i) The schematic illustrates the preparation of decellularized collagen, showing two strips cut in both parallel and perpendicular directions. (j) Mechanical properties of the collagen matrices measured by dynamic mechanical analysis. Double-sided arrows denote the alignment directions. (k) The schematic illustrates the procedure for force analysis, where cell-cell contraction compacts the matrix.

Next, we characterize the microstructure of the collagen ECM. To visualize the microstructure without the encapsulated cells, we add Triton X-100, which disrupts the phospholipid membrane and decellularizes the collagen matrix. Using scanning electron microscopy, we find that cells modify the collagen microstructure to align along the direction imposed by the PEG hydrogels. The initially isotropic collagen (Fig. 3g) develops increased orientational anisotropy through its interaction with HDFs. Furthermore, the structures also densify, resulting in thicker bundles (Fig. 3h).

We anticipate that the initially isotropic collagen will also develop increased mechanical anisotropy through alignment induced by cellular activities. To assess the mechanical properties of the remodeled matrix in both the parallel (∥) and orthogonal (⊥) directions, strips are cut from the decellularized matrix along these two orientations (Fig. 3i). Tensile testing is carried out using dynamic mechanical analysis (Fig. S6). To overcome the softness of collagen (∼ kPa), which prevents it from being a self-supporting gel outside of an aqueous environment, we place an elastomer sheet (∼ 100 kPa) underneath it as a support. By measuring stress-strain curves (Fig. S6) of the elastomer alone versus elastomer plus collagen, we find the modulus of the collagen by taking the difference between these two measurements. This way, we obtain moduli *G*_∥_ = 8.6 kPa, and *G*_⊥_ = 6.6 kPa for collagen gel that undergoes remodeling by aligned cells after 5 days, compared to *G* = 2.20 kPa for an isotropic collagen gel on Day 0 (Fig. 3j). *G*_∥_ and *G*_⊥_ both increase after several days of cell-culturing, which signifies that cells compress the matrix, making it denser and stiffer. This effect is more pronounced along the alignment direction. These observations complement the microstructure characterization in Fig. 3g,h.

The alignment of cells results in an anisotropic contraction, even though the mechanical property is stiffer along that direction. Based on cell positions tracked in Fig. 3f, we construct a simple model that quantitatively accounts for the varying rate of collagen contraction (Fig. 3k). Here, we make several simplifying assumptions: 1) The cell distribution snapshot represents the distribution throughout the entire contraction process. 2) On the scale of the matrix, the matrix properties can be treated as uniform along the two orthogonal directions. 3) The contractile force can be entirely attributed to the nearest neighbor, with each pair exerting the same average force 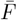 along angle *θ*_*ij*_, which refers to the angle formed between the center-to-center line connecting cells *i* and *j* and the alignment direction (Fig. 3k, zoomed-in view). The total force along the alignment direction is therefore 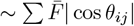, whereas the total force along the perpendicular direction is 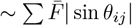 As a result, the theoretically derived rate of contraction along the alignment direction to that along the perpendicular direction is :

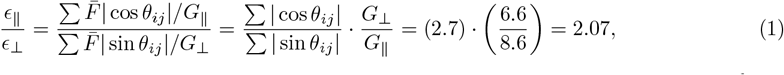

which is consistent with the observed anisotropic rate of contraction across multiple samples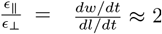, found in Fig. 2e.

### 2.4 Patterning cell alignment for traction-mediated shape changes

Replicating PEG fiber patterns in cell-laden matrices opens new avenues beyond uniaxial contraction. By establishing a spatially inhomogeneous alignment field, we could potentially realize spontaneous, programmable, shape changes — a process we refer to as *in vitro* “morphogenesis”. This calls for defining not only the directionality but also the specific locations where oriented cell forces are applied. Achieving this requires a resolution beyond what is currently achievable by 3D printing, which has a minimum resolution of 100s of *µm*. In contrast, photopatterning is a well-established technique for programming order in LCEs [48] and LCs [49] that can achieve resolutions of 10s of *µm*. Here, a third method, photopatterning, is used to create spatially heterogeneous PEG fibrous hydrogels. This pattern was then translated into HDF ordering using a sequential deposition approach.

We create a spatially heterogeneous commanding field by using a custom-built photopatterning setup (Fig. 4a) following [50]. Briefly, we first coat a glass slide with an azobenzene dye, and illuminate it with polarized light through a mask, region by region. Afterward, we spin-coat a layer of liquid crystal monomer RM257, and then UV crosslink the RM257. Two such slides are assembled under the microscope with their patterns aligned and facing each other. When DSCG-PEGDA hydrogel precursor solution is injected, it follows the alignment direction (also known as the “director field”) of the RM257 layer (Fig. 4b). An example of a spatially inhomogeneous field is one with topological defects, where the order cannot be defined (red dots). Defects can be classified based on their charges, or winding number, referring to the total number of times the director field rotates around this point [51]. A -1 defect is illustrated in Fig. 4c and is created by a triangular slit (outlined in blue). As one goes around the center of the defect by 2*π* in a clockwise fashion, the polarizer, which determines the director, undergoes a -2*π* rotation synchronously. The ratio of their rotation defines topological charge, which is 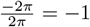 in this case. If we reduce the rotational speed of the polarizer by half, we obtain a 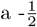 defect, also known as a trefoil, as shown in Fig. 4d. If instead, the polarizer rotates in the same direction as the slit, we obtain a radial +1 defect, as shown in Fig. 4e. If the polarizer remains perpendicular to the slit orientation, then a circular +1 defect can be obtained, as shown in Fig. 4f. We also present polarized microscopy images in (ii), and visualization of the hydrogel fibers by phase contrast mode in (iii). A *λ*-plate is used to distinguish between positive and negative deviations from zero. As a result, fibers with an orientation of 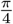 appear blue, while those with an orientation of 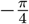 appear yellow. In Fig. S7, we provide zoomed-in views of phase contrast images to show the resolutions are around 10 *µ*m or less. The photopatterning setup provides a unique opportunity to program the orientation of the hydrogel at fine resolution into predefined patterns, as shown in Fig. 4c-g. Replicating such a system requires minimal equipment and no synthesis, only commercially available chemicals, making it highly adaptable for use across soft matter and biological laboratories.

**Figure 4:**
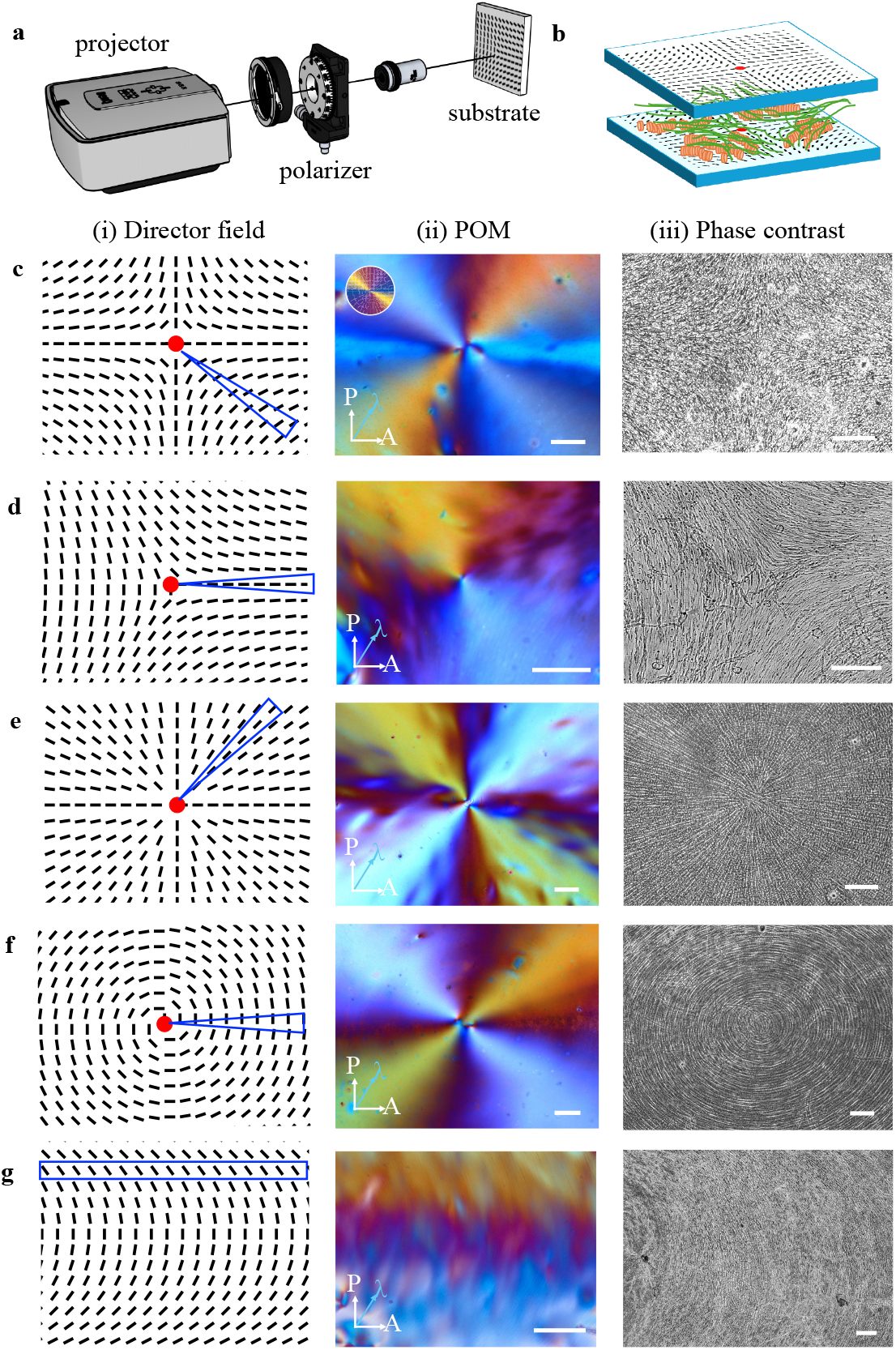
Creating spatially defined orientational field by photopatterning. (a) Schematics of the projector display setup. (b) Schematics for preparation of patterned hydrogels. DSCG follows the alignment imposed by the SD1 dye molecules at the substrate. The structure is fixed by crosslinking the PEGDA (not shown). (c-g) A variety of spatially varying structures are generated. (i) Schematics of the director field, where the red dots denote the defect. The blue rectangle and triangle indicate the slit, through which a region of the substrate is illuminated with a fixed polarizer orientation. After the entire region has been patterned, the final patterns are shown in (ii) Polarized optical micrographs (POM). A *λ* plate is inserted to disambiguate orientation greater than or less than *π*, represented by the color wheel. (iii) Phase contrast micrographs of the crosslinked structure of the patterned hydrogels. All scale bars are 100 *µ*m.

Compared to half-integer defects such as the one shown in Fig. 4d, integer defects, such as the one shown in Fig. 4c,e,f, are far less common in 2D monolayers. They can only be stabilized under confinement [52] or with strong boundary guidance [26]. Nevertheless, radial cell configurations naturally occur at tumor sites [53] and in nervous systems [54], while circular arrangements are also commonly observed in cross-sections of blood vessels [55]. Therefore, replicating these morphologies is a valuable target for tissue engineering. We illustrate the ability to pattern hydrogels of both radial (Fig. 4e) and circular (Fig. 4f) morphologies. Introducing these defects through surface alignment and extending them into a 3D volume opens new avenues for supracellular assembly with realistic morphologies. Finally, we illustrate that the accessible patterns are not restricted to those with rotational symmetry, as illustrated by the “C” shaped configuration in Fig. 4g. By selectively illuminating regions of the substrate while simultaneously rotating the polarization, non-rotationally symmetric fields can also be achieved.

As illustrated in Fig. 2, directional cell alignment can result in varying contraction rates along and perpendicular to the alignment axis. Could this suggest that patterned cell assembly might also induce specific macroscopic shape changes? By depositing a cell-collagen mixture on top of the hydrogel fibers in a similar fashion, and after allowing the cells to grow for 3-5 days in contact with the photopatterned hydrogel, we find that HDFs also replicate the hydrogel pattern underneath. Three such configurations are shown in Fig. 5a-c, which correspond to the director field configuration in Fig. 4c-e. Once the cells have grown to a density of 800 mm^−2^, we wash away excess cells not encapsulated in the collagen matrix by adding Trypsin-EDTA, which also helps dissociate the attached cells at the boundary of the collagen from the hydrogel surface, detaching the collagen matrix from the hydrogel substrate. Immediately afterwards, cells within the collagen round up but remain encapsulated in the matrix. We then incubate the cell-laden matrix in complete media to allow them to recover. When they re-extend after several hours, they do so in a manner that follows the previous pattern, as the cells have already remodeled the collagen by this point, as shown in Fig. 3h.

**Figure 5:**
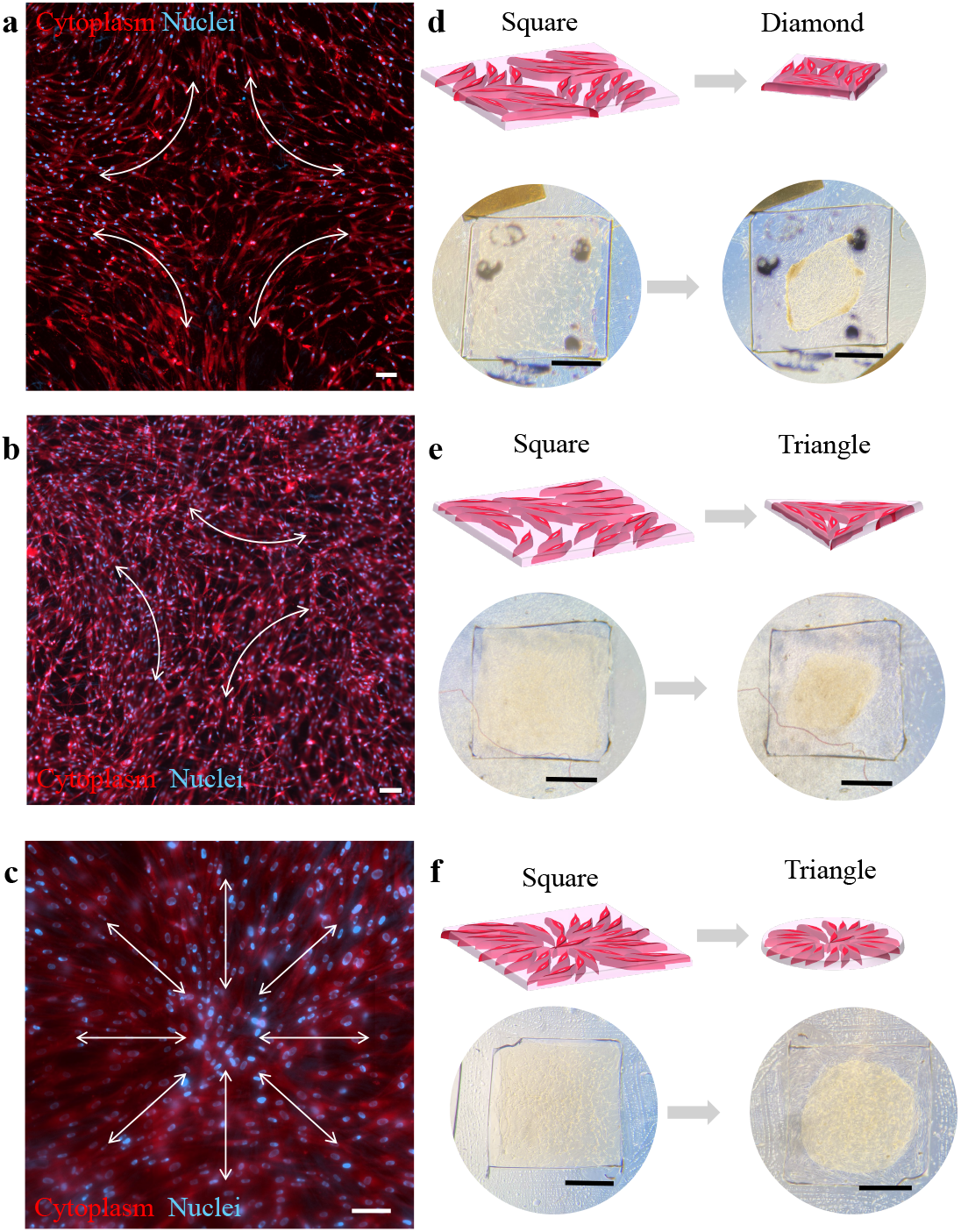
Programmable shape transformations. (a-c) Fluorescent micrographs of encapsulated cells in hydrogels, arranged in the patterns of topological defects of charge (a) -1, (b) 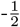, and (c) radial +1. The cytoplasm and nucleus are visualized using CellTracker DeepRed and DAPI, respectively. White double-sided arrows help visualize the direction of collective cell forces. (d-f) Corresponding schematic of cell contraction, followed by images of macroscopic contraction of the collagen sheets. The image on the left shows the collagen sheet on day 1, while the images on the right show the sheet after contraction. The scale bars are 100 *µ*m in panels (a)-(c) and 1 mm in panels (d)-(f). The region within the collagen gels is highlighted for clarity.

In the subsequent 24 hours, the cell-laden collagen matrices undergo shape changes due to cell traction following their orientation and collective forces (denoted by double-sided arrows in Fig. 5a-c). For instance, the collective alignment of cells around a defect with charge -1 generates a field analogous to a force quadrupole (Fig. 5a), causing the collagen matrix to transition from a square shape to a diamond (Fig. 5d). An analogous scenario arises for cell alignment following a pattern of the topological charge Of 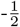 (Fig. 5b), where collective cell forces induce the transformation from the square to a triangle (Fig. 5e). Finally, we demonstrate that a radially oriented cell field (Fig. 5c) can transform a square into a circle (Fig. 5f). Although all three hydrogels started with the same square shape, the difference in the orientation of the encapsulated cells and their collective forces resulted in distinct final shapes for each scenario. We highlight the region of the collagen for clarification, Unhighlighted version and additional instances of programmable shape changes driven by these orientation fields are shown in Fig. S8-10. Combining different shapes is expected to allow for more complex shape transformations. This straightforward *in vitro* system highlights how collective alignment may drive programmable shape changes.

## 3. Conclusion

In this work, we combine the biocompatibility of PEG hydrogels with the biodegradability of collagen to engineer supracellular assemblies. Our approach is inspired by the literature on liquid crystalline elastomers, where mesogenic molecules are organized directionally to generate anisotropic forces. In our system, however, the contractile forces are spontaneously generated by the cells. We first characterize the anisotropic structure of PEG hydrogels, which is the source of anisotropy. Subsequently, we show that the hydrogel can guide the uniaxial alignment of cells encapsulated in collagen, in contact with the hydrogel. These cells modify the collagen matrix they inhabit. The tracked anisotropic contraction rate is in good agreement with the one calculated from the observed cell orientation distribution and measured mechanical properties.

By precisely controlling the morphology and orientation of the PEG hydrogel through photopatterning, we successfully replicate this organization amongst the encapsulated cells. This approach allowed us to program the collective forces exerted by the cells, resulting in predefined macroscopic shape changes in the cell-laden matrix. Our work supports the argument that an appropriately designed initial state can initiate anisotropic, collective cell proliferation and remodeling, culminating in mechanical stress that deforms the matrix. Our technique enables precise control of collective cell orientation. We show that by programming the initial state, highly anisotropic and coupled bio-chemo-mechanical dynamics can be captured *in vitro*.

While we demonstrate our method using pure collagen, the technique is expected to be generalizable to other degradable matrices [56, 57]. We anticipate that the method in this manuscript can be combined with techniques like microstamping [58] and lithography [59] to fully define shape transformation. With the versatility of the patterning approach, our platform can be adopted by a variety of biological laboratories, providing a reliable and cost-effective setting for assessing the dynamic hydrogel-based matrices, and paving the way for potential applications ranging from wound dressings, hygiene products, tissue regeneration, to disease models.

## 4 Experimental Section

### 4.1 Materials

The monomer poly(ethylene glycol) diacrylate (average Mn = 250, PEGDA250), lyotropic liquid crystal disodium cromoglycate (DSCG), photoinitiator lithium phenyl-2,4,6-trimethylbenzoylphosphinate (LAP), azo polymerization initiator azobisisobutyronitrile (AIBN), sodium dodecyl sulfate (SDS), Triton X-100, N,N-dimethylformamide (DMF), acetone, isopropanol were purchased from Sigma-Aldrich (St. Louis, MO), and used without modification. Sylgard 184 polydimethylsiloxane (PDMS) Elastomer Kit was purchased from Dow Chemical (Midland, MI). Azodye SD1 dye was purchased from DIC Corporation (Tokyo, Japan). Irgacure-651 was purchased from MedKoo Biosciences (Durham, NC). Water was dispensed from a Milli-Q system with resistivity = 18.2 MΩ·cm at 25 °C.

Human dermal fibroblast (HDF, neonatal) cells were purchased from the American Type Culture Collection (Manassas, VA, catalog no. PCS-201-010). Dulbecco’s modified Eagle’s Medium (DMEM), fetal bovine serum (FBS), phosphate-buffered saline (PBS), Trypsin-EDTA (0.25%), and penicillin-streptomycin (pen-strep) were purchased from Gibco (Grand Island, NY). Paraformaldehyde (PFA), Cell-Tracker Far-Red (C34552), and Hoechst 33342 (H3570) were purchased from ThermoFisher (Waltham, MA). High concentration, rat tail, type I collagen (10.25 mg/mL) was purchased from Corning Inc (Corning, NY). RhodamineB-PEG-SH was purchased from Biopharma PEG (MW = 2k, Watertown, MA).

### 4.2 Fabricating hydrogel substrates

All glass slides and coverslips were thoroughly cleaned with detergent in a sonication bath, then rinsed with acetone and isopropanol, followed by plasma cleaning before use.

Uniformly aligned hydrogel fibers were generated following the procedure in [40]. Briefly, a solution of 5% PEGDA250, 15% DSCG, and 1% LAP was introduced into a glass chamber. The glass chamber was made by sand-paper rubbed glass slides, with rubbed sides facing each other, separated by 20 *µ*m bead spacers Afterward, the hydrogel precursor solution was injected into the chamber via capillary force, and heated to the isotropic phase. The filled-in chambers were slowly cooled down to the nematic phase (T = 5 °C), maintained at that temperature on a Linkam thermal stage (The McCrone Group, Westmont, IL), and crosslinked for 2 hours by an ultraviolet lamp (*λ* = 365 nm, power = 6 W). Thereafter, the chamber was opened with a razor blade and immersed in water to remove the DSCG.

### 4.3 Photopatterning of the hydrogel

Patterned lyotropic alignments were prepared by photopatterning using procedures detailed in [60, 49]. Briefly, a solution of 0.2 wt% SD1 in DMF was evenly deposited on the glass slide by spin coating at 3000 rpm for 30 seconds. The coated glasses were illuminated by a projection display, protected from the light, and a thin layer (7 wt% RM257, 0.05 wt% Irgacure 651 in toluene) was deposited on the surface to stabilize the dye [46]. The patterned glass slides were assembled under the microscope, into a chamber with coated sides facing each other, separated by spacer beads (2*a* = 4 - 4.5 *µ*m, Spherotech Inc, Lake Forest, IL). The hydrogel precursor solution was filled in, crosslinked, and rinsed, as described in Section 4.2.

### 4.4 X-ray Diffraction

Scattering samples were prepared by injecting a solution of 5% PEGDA250, 15% DSCG, and 1% LAP using a 22 Gauge needle at the nematic phase into a 1.5 mm quartz tube (Charles Supper Company Inc, Westborough, MA), at 1 ∼ mm/s. The quartz tube was placed in the freezer for 20 minutes beforehand so as not to raise the temperature of the PEGDA-DSCG solution during loading. The tube was sealed by parafilm and the sample was crosslinked with a long wavelength (365 nm, 6W) lamp for 2 hours at T = 5 °C. To rinse off the DSCG, the quartz tube was immersed in 50 mL of DI water overnight to ensure DSCG diffused away. The sample was scattered using a custom-built, high-brilliance, liquid gallium X-ray source located in UCSB, each for 2 hours. The wide-angle X-ray scattering (WAXS) and small-angle X-ray scattering (SAXS) data were reduced by IGOR Pro (WaveMetrics Inc). The background was subtracted from the raw data.

### 4.5 Cell culture and handling

The HDF cells were maintained at 37°C with 5% CO_2_ in complete media: DMEM supplemented with 10 vol% FBS and 1x pen-strep. The media was refreshed every 2 days. Whenever the cells reached 70-80% confluency, they were lifted by adding Trypsin-EDTA, centrifuged, and resuspended at 20-30% density. Early passages (less than 20) of HDFs were used in the study. No significant difference in cell behaviors between passages was noted.

### 4.6 Collagen sheet fabrication

The spacer that defines the outer shape of the collagen matrix was cut from a crosslinked 100 *µ*m PDMS sheet. To make the PDMS spacer, the base and the curing agent were mixed in a 10:1 ratio, degassed under vacuum to remove bubbles, and cured at 80 °C for 2 hours. The PDMS spacer was bonded to the hydrogel substrates (as discussed above) to form a mold, into which a mixture of neutralized collagen and cells was added.

A total of 50 *µL* of neutralized collagen solution, consisting of 1 part 10x PBS, 4 parts DI water, and 5 parts acid-solubilized collagen (concentration = 10.25 mg/mL), was mixed on ice. A dense suspension of HDFs (cell density ∼ 1.6 × 10^6^ cells/mL) was subsequently combined with the neutralized collagen, resulting in a final cell concentration of ∼ 8 × 10^5^ cells/mL, and a final collagen concentration of 4 mg/mL. Additional spacer beads (diameter 2a = 30-40 *µ*m) were added on top of the PDMS spacer prior to placing another coverslip to flatten the collagen. The additional gap created by the spacer beads also facilitated media diffusion into the hydrogel. The collagen matrix (total thickness ∼ 130 *µ*m) was crosslinked at 37 °C in the incubator for 1 hour, then more complete media was added. After cells were grown on the hydrogel substrate for 5 days, the collagen was released from the collagen fiber, resulting in a free-standing sheet with programmable cell orientation.

### 4.7 Decellularization

To destabilize the cell membrane, 0.1 wt% Triton X-100 in PBS was added to the cell-laden matrix, and the matrix was incubated at room temperature for 5 minutes. Finally, the collagen matrix was rinsed with PBS three times and maintained in PBS.

### 4.8 Microscopy techniques

To visualize the hydrogel, PEGDA250 was fluorescently tagged by reacting by mixing it with a solution of 10 wt% PEGDA250, 1 wt% rhodamineB-PEG-SH, and 1 wt% AIBN in DMSO, in a vacuum oven at 85°C for 1 hour. Hydrogel fibers were labeled by mixing tagged: untagged PEGDA250 in a 1:10 ratio, and the hydrogel was fabricated as before. HDF cells in collagen were permeated by 4% PFA, and stained by CellTracker and Hoechst 33342 following the manufacturer’s procedures. Imaging was performed using a Zeiss Axio Observer 7 microscope outfitted with a computer-controlled motorized sample stage, motorized auto-focus objectives, and an Axiocam 702 monochromatic camera. A standard processing module in the Zeiss ZEN software was used to stitch together multiple fields of view. To image 3D cellular structure, confocal microscopy was performed using a Zeiss LSM 880 Confocal Microscope.

### 4.9 Scanning electron microscopy

Hydrogel and collagen samples of approximately 1 cm × 1 cm in size are affixed to a holder using double-sided adhesive carbon tapes and sputtered with a thin layer of iridium to prevent sample charging problems. Samples were recorded with a typical accelerating voltage of 5 kV, using a Hitachi SU8230 UHR Cold Field Emission (CFE) Scanning Electron Microscope.

### 4.10 Mechanical testing

Collagen marix, produced without cells, or decellularized after cell growth, were measured using dynamic mechanical analysis along both the alignment and orthogonal directions. To circumvent the softness of collagen (∼ kPa), which makes it difficult to clamp onto the holder, we reinforce it with an elastomer sheet, which was made with a 40:1 weight ratio of prepolymer base to curing agent using the PDMS kit according to [61], with a modulus of ∼ 100 kPa. Due to batch-to-batch variation, the mechanical properties of each batch of PDMS backing are re-measured. By measuring stress-strain curves of the elastomer alone versus elastomer plus collagen, we found the modulus of the collagen by taking the difference between these two measurements.

## Conflict of interest

The authors declare no conflict of interest.

## Supporting Information

Supporting Information is available from the Wiley Online Library.

## Data Availability Statement

The data collected and the analysis codes generated for this study will be made available through a permanent DOI on Dryad upon acceptance of this manuscript.

## Acknowledgements

We thank Joe Wolenski for training in using the confocal microscope, Amit Datye and David Gaetano for their assistance with the dynamic mechanical analysis, and Phillip Kohl and Youli Li for their help with the X-ray scattering. The authors acknowledge the use of experimental facilities at Yale Institute for Nanoscience and Quantum Engineering, Yale Science Hill Imaging Facility, the Mechanical and Thermal Analysis Instrumentation Core, and the scattering facility at UCSB funded by the BioPACIFIC Materials Innovation Platform of the National Science Foundation under Grant No. DMR-1933487 (NSF BioPACIFIC MIP). Y.L. acknowledges the support by the National Science Foundation under Grant No. OAC-2411044.

## Notes

### Competing Interest Statement

The authors have declared no competing interest.

