## Supplemental Figure S1-S10 for "Cell-sheet shape transformation by internally-driven, oriented forces"

### Supporting Information for “Cell-sheet shape transformation by internally-driven, oriented forces”

November 28, 2024

#### Additional Figures

Figure S1. Scanning electron microscopy of the dried PEG hydrogel microstructure prepared by flow, and photopatterning.

Figure S2. Anisotropic microstructure of the PEG hydrogels.

Figure S3. Additional scattering results: Scattering measurements of PEG hydrogels containing or excluding DSCG.

Figure S4. Control experiment showing that the cells in collagen matrix fail to achieve alignment when they are grown on hydrophobic liquid crystal elastomer fibers.

Figure S5. Custom analysis procedure for finding the dimension of the collagen sheet.

Figure S6. Tensile strength measurements for contracting cell-modified hydrogels.

Figure S7. Resolution of photopatterned defect hydrogel fibers.

Figure S8-S10. Additional results for shape transformation of cell-laden collagen sheets.

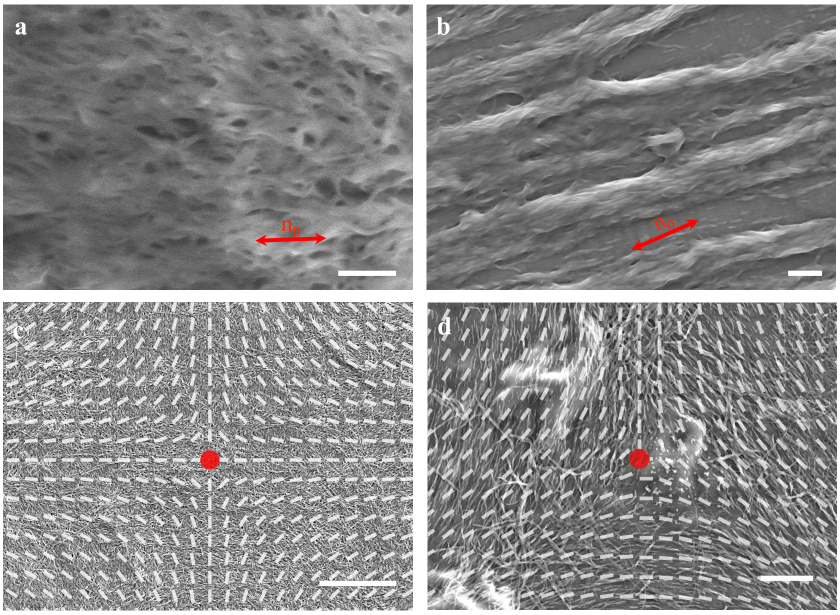


Figure S1: **Scanning electron microscopy of the dried PEG hydrogel microstructures prepared by different methods.** Micrographs of the structures prepared by flow alignments in (a) and photoalignment substrates in (b-d). The patterns are unidirectional alignment in (b), zoomed-in -1 defect in (c), and -$\frac{1}{2}$ defect in (d). All images display the fibrous morphology of the PEG hydrogels. The scale bar is 0.5 $\mu$m in (a), 1 $\mu$m in (b), 20 $\mu$m in (c), and 5 $\mu$m in (d). Red double-sided arrows denote the imposed alignment direction. Red dots denote the center of the defect. The photopatterned director field is shown as an overlay. The formation of the fiber is driven by a reaction-diffusion process, so the microstructure remains largely consistent, regardless of the alignment mechanism used.

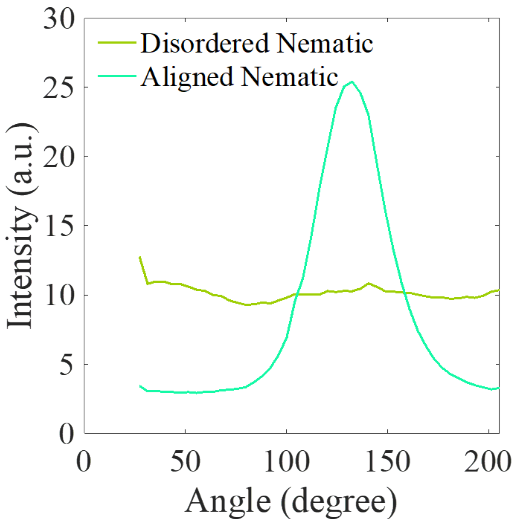


Figure S2: **Anisotropic microstructure of the PEG hydrogels**. Integrated wide-angle x-ray scattering intensities over angle for anisotropic and disordered PEG hydrogel fibers crosslinked at nematic phase (15 wt% DSCG and 5 wt% PEGDA). The curve of anisotropic PEG hydrogel fiber shows one significant peak, while the curve of the disordered PEG hydrogel fiber stays flat.

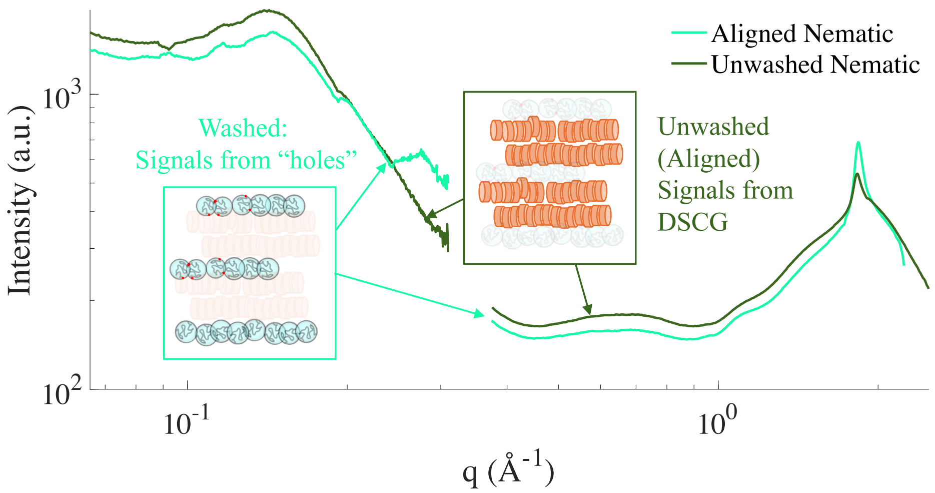


Figure S3: **Scattering measurements of PEG hydrogels containing or excluding DSCG**. To understand the origin of the signal, we perform X-ray scattering before and after washing the DSCG. Combined SAXS and WAXS scattering intensities for washed and unwashed PEG fibers crosslinked at nematic phase (15 wt% DSCG and 5 wt% PEGDA). The insets show schematics of the structural components that scatter the strongest signals. After rinsing out the DSCG, the scattering signal remains largely unchanged.

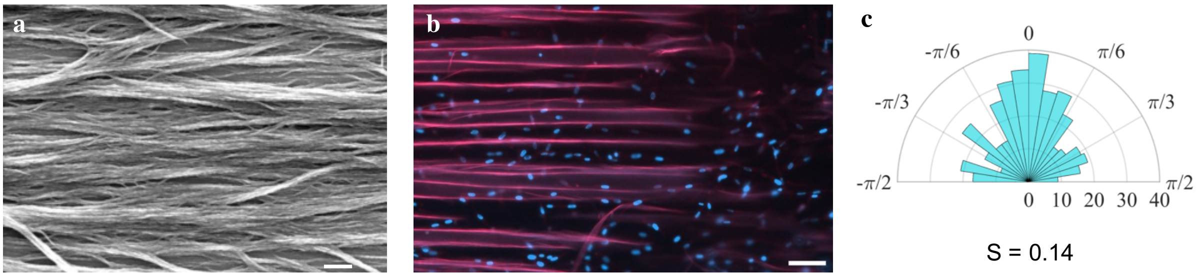


Figure S4: **Abscence of cell alignment when they are grown on hydrophobic LCE fibers**. (a) Scanning electron microscopy of uniform LCE fiber. The scale bar is 1 $\mu$m. (b) HDF in collagen cultured on hydrophobic LCE fibers. Cell nuclei are stained by Hoechst. The scale bar is 100 $\mu$m. (c) Polar histogram of the nucleus orientation on LCE fibers. The LCE fibers were prepared using a method that is nearly identical to the PEG hydrogel fabrication. However, in place of the hydrogel precursor solution, a solution of 7 wt% reactive monomer RM257, 92 wt% small molecule liquid crystal mixture, and 1 wt% photoinitiator was introduced into the pre-assembled chamber at isotropic temperature T = 120 °C, and slowly cooled down to nematic phase T = 20 °C. The sample was cured under UV light with power 6 mW/cm$^2$ at 365 nm wavelength for 1 h to crosslink the reactive monomer completely. Afterward, the samples were immersed in hexane overnight to wash away the uncrosslinked small molecule liquid crystals. Finally, the samples were dipped in liquid nitrogen for a few seconds and split with a razor blade. As a result, stiff (100s of MPa) LCE fibers were generated instead of soft hydrogels (10s of Pa).

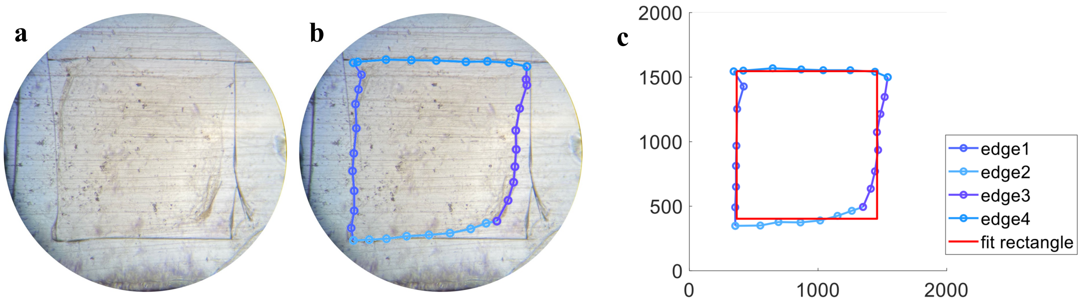


Figure S5: **Custom analysis procedure for finding the dimension of the collagen sheet.** (a) Original image. (b) The edge of the cell-laden matrix was traced manually, each edge is described by 5-7 points. We then compute the average of the vertical and horizontal edges to fit a rectangle to find the width $w$ and length $l$. (c) The red solid line denotes the fitted rectangle.

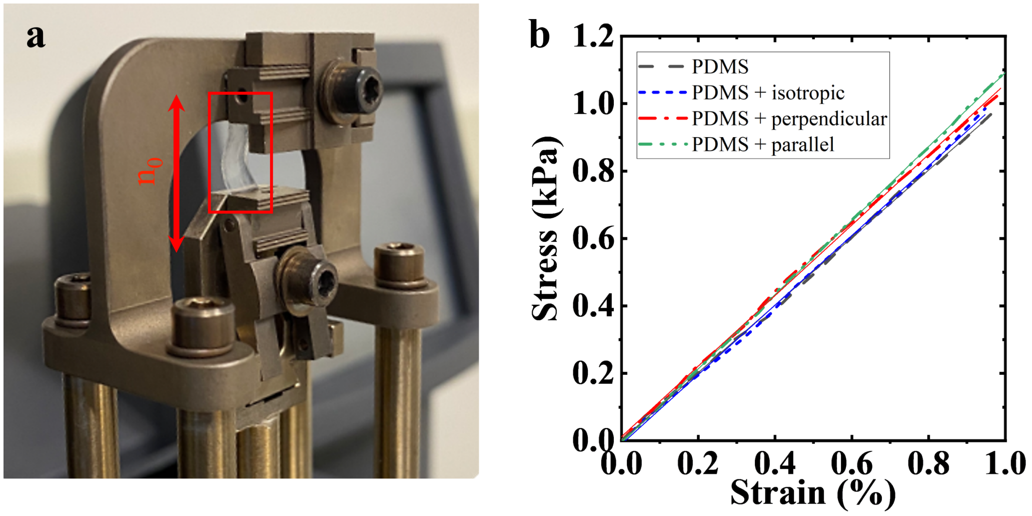


Figure S6: **Tensile strength measurements for cell-modified hydrogels.**(a) Dynamic mechanical analysis setup. The PDMS and the hydrogel were cut to identical size, so upon stretching, both undergo identical strain $\epsilon_1 = \epsilon_2 = \epsilon$. (b) Stress-stress curves for various preparations of the hydrogel and the PDMS backing, which is used to support the hydrogels.

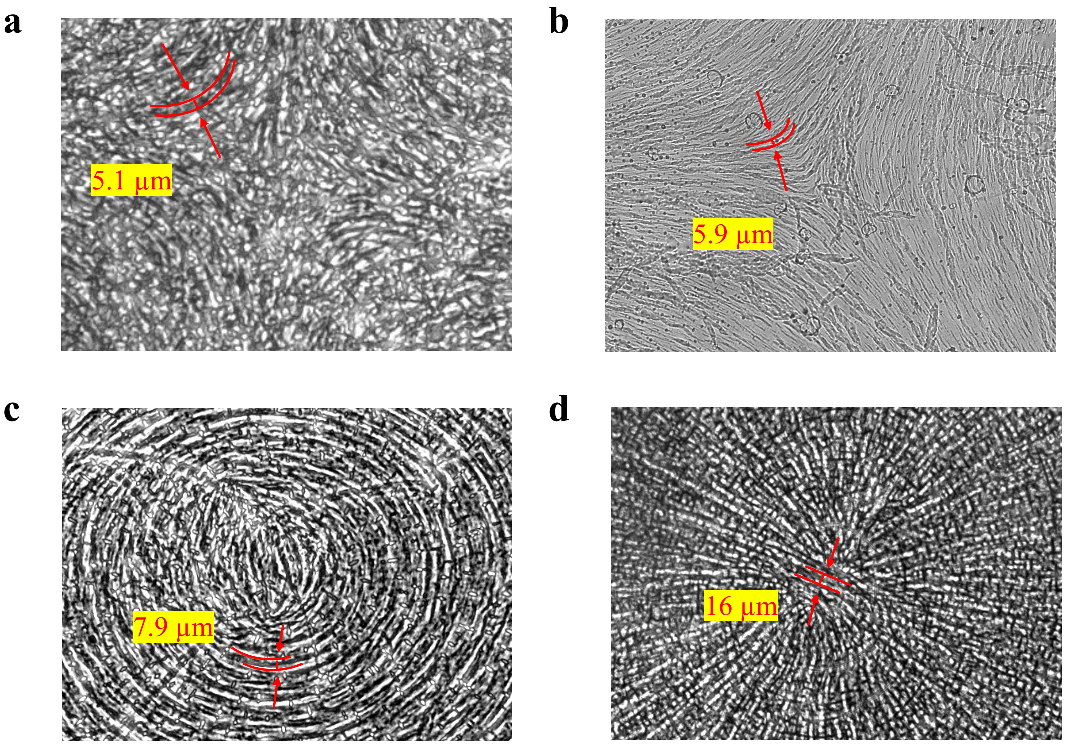


Figure S7: **The resolution of photopatterned defect hydrogels.** Red lines and arrows show the distance between two smallest resolvable grooves, which we define as the resolution. The number shows the resolution in each image.

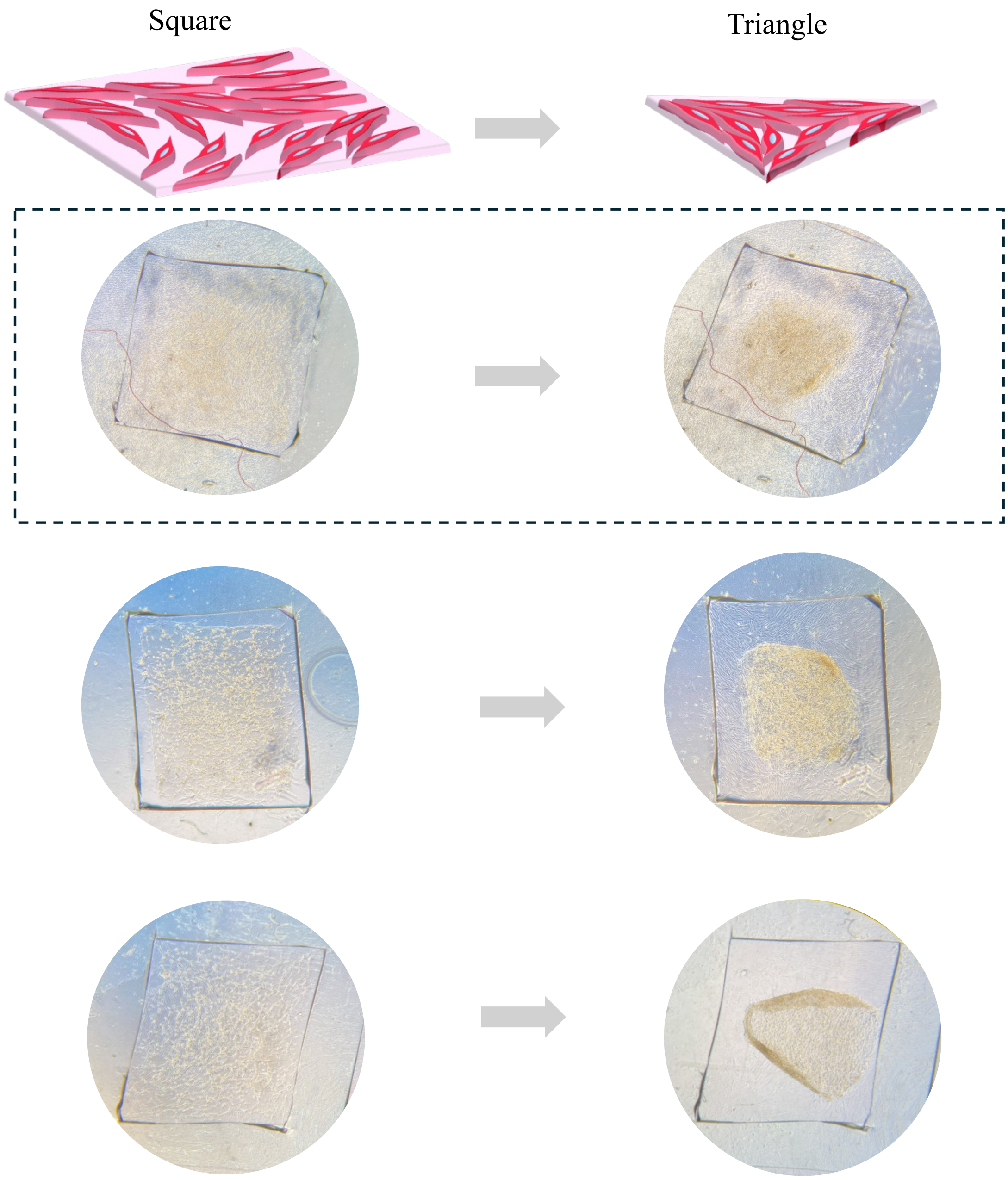


Figure S8: **Additional results for shape transformation of cell-laden collagen sheet from a square to a triangle.** The cells are arranged in a -1/2 topological defect pattern. The row boxed with dotted lines corresponds to the figure in the main text, without highlighting the collagen matrix region.

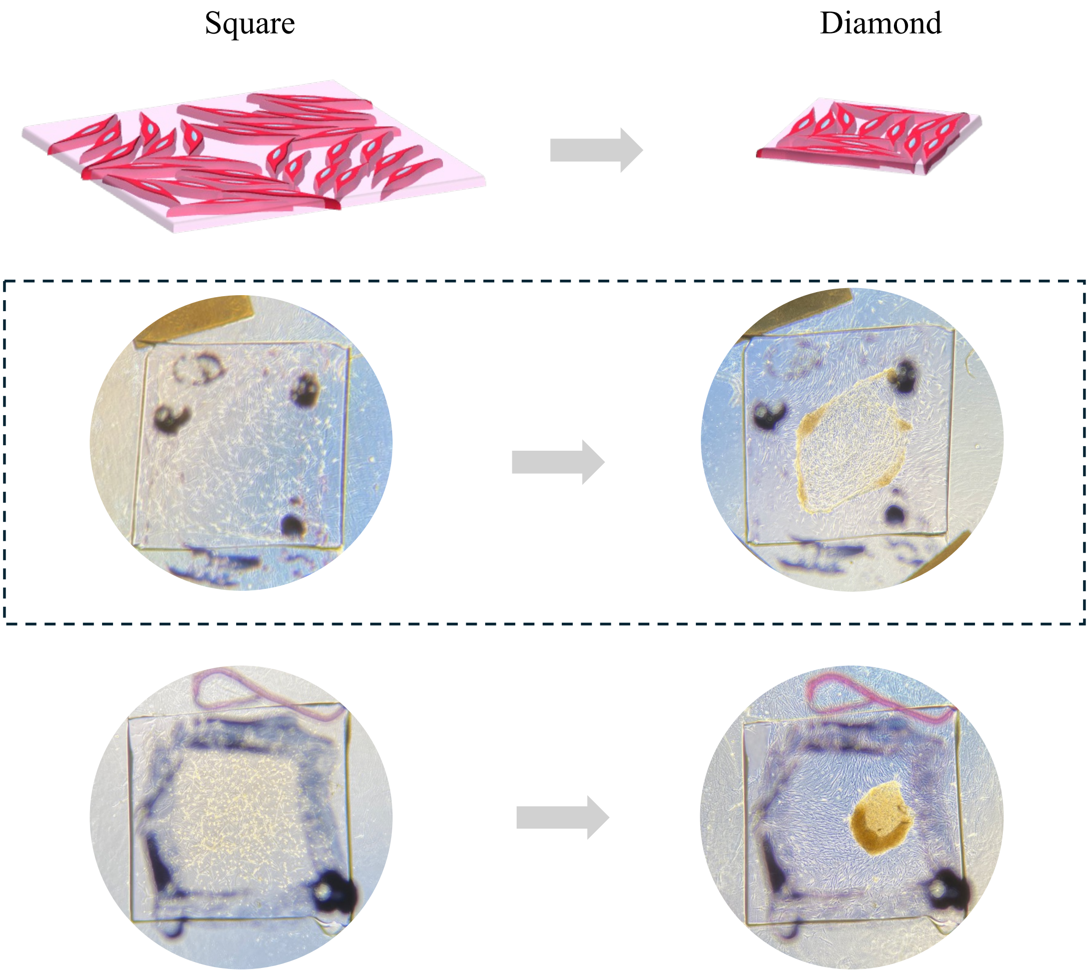


Figure S9: **Additional results for shape transformation of cell-laden collagen sheet from a square to a diamond.** The cells are arranged in a -1 topological defect pattern. The row boxed with dotted lines corresponds to the figure in the main text, without highlighting the collagen matrix region.

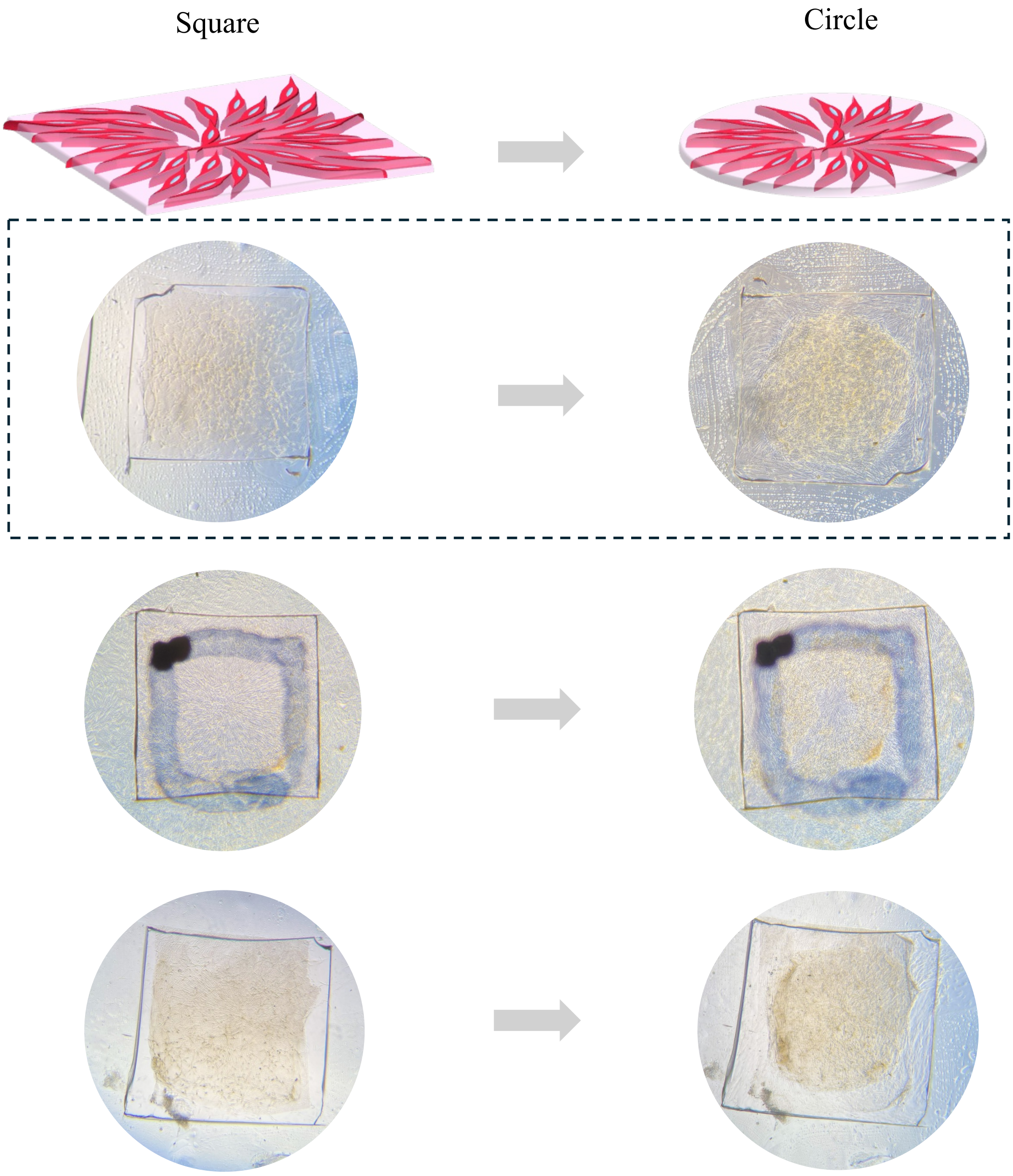


Figure S10: **Additional results for shape transformation of cell-laden collagen sheet from a square to a circle.** The cells are arranged in a radial +1 topological defect pattern. The row boxed with dotted lines corresponds to the figure in the main text, without highlighting the collagen matrix region.
